# A receptor-like kinase mediated phosphorylation of Gα protein affects signaling during nodulation

**DOI:** 10.1101/2019.12.11.873190

**Authors:** Swarup Roy Choudhury, Sona Pandey

## Abstract

- Heterotrimeric G-proteins, comprised of Gα, Gβ and Gγ subunits regulate signaling in eukaryotes. In metazoans, G-proteins are activated by GPCR-mediated GDP to GTP exchange on Gα; however, the role of receptors in regulating plant G-protein signaling remains equivocal. Mounting evidence points to the involvement of receptor-like kinases (RLKs) in regulating plant G-protein signaling pathways, but their mechanistic details remain limited. We have previously shown that during soybean nodulation, the nod factor receptor 1 (NFR1) interacts with G-protein components and indirectly controls signaling.
- We explored the direct regulation of G-protein signaling by RLKs using protein-protein interactions, receptor-mediated phosphorylation and the effects of such phosphorylations on soybean nodule formation.
- Results presented in this study demonstrate a direct, phosphorylation-based regulation of Gα by symbiosis receptor kinase (SymRK). SymRKs interact with and phosphorylate Gα at multiple residues, including two in its active site, which abolishes GTP binding. In addition, phospho-mimetic Gα fails to interact with Gβγ, potentially allowing for constitutive signaling by the freed Gβγ.
- These results uncover a novel mechanism of G-protein cycle regulation in plants where receptor-mediated phosphorylation of Gα not only affects its activity, but also influences the availability of its signaling partners, thereby exerting a two-pronged control on signaling.

## INTRODUCTION

Heterotrimeric G-proteins comprised of Gα, Gβ and Gγ proteins are key signaling components in eukaryotes. As per the classical paradigm, when Gα is GDP-bound, the proteins maintain a trimeric conformation and are inactive. In this form, the proteins are associated with a cognate G-protein coupled receptor (GPCR). Ligand binding to the GPCR causes a change in its conformation, which allows exchange of GTP for GDP on Gα. GTP-bound Gα dissociates from the Gβγ dimer and both these entities can interact with downstream effectors to transduce the signal. This represents the active signaling status of G-proteins (Offermanns, 2003). The inherent GTPase activity of Gα, in conjunction with GTPase activity accelerating proteins (GAPs) such as the regulator of G-protein signaling (RGS) proteins regenerate GDP-bound Gα, which re-associates with the Gβγ dimer to form the inactive heterotrimer, ready for the next cycle of activation (Offermanns, 2003; McCudden *et al.*, 2005). Depending on the pathway, the signal may primarily be transduced by the active Gα protein, with the role of Gβγ only to sequester it in its inactive form. Alternatively, the signal may be transduced by the activity of both Gα and Gβγ proteins. In some cases, the Gβγ proteins function as the main signal transducers, and require the Gα activation only to be freed from the complex. Experimental evidence exists for each of these scenarios (Koelle, 2006; Dupre *et al.*, 2009; Pandey *et al.*, 2010).

Similar to the metazoan systems, the plant Gα proteins are active when GTP-bound, and exhibit guanine nucleotide-dependent monomeric or trimeric status, but the extent to which a classic GPCR is required for their activation is unclear (Urano *et al.*, 2012; Urano & Jones, 2014; Roy Choudhury & Pandey, 2016). The proteins do interact with many GPCR-like plant proteins (Pandey & Assmann, 2004; Gookin *et al.*, 2008; Pandey *et al.*, 2009; Roy Choudhury & Pandey, 2016), but there is no evidence yet for their role in facilitating GDP to GTP exchange (Pandey, 2019).

Recent work suggests that in plants, Gα proteins also interact with the highly prevalent receptor-like kinases (RLKs) (Bommert *et al.*, 2013; Liu *et al.*, 2013; Ishida *et al.*, 2014; Aranda-Sicilia *et al.*, 2015; Liang *et al.*, 2016; Trusov & Botella, 2016; Liang *et al.*, 2018; Pandey & Vijayakumar, 2018; Peng *et al.*, 2018; Yu *et al.*, 2018; Pandey, 2019). RLKs have one to three transmembrane domains that connect the extracellular region to the intracellular kinase domain (Shiu & Bleecker, 2001). They typically function as multi-protein complexes comprising of one or more RLK, additional receptor-like protein (RLP) and/or cytosolic proteins. RLKs typically act by phosphorylating their associated co-receptors, RLPs or their downstream targets (*De Smet et al., 2009*). Some of the best characterized examples of such RLK complexes include the BRI1-BAK1 receptor pair and the FLS2-BAK1 receptor pair, where growth and defense responses of plants are mediated by distinct RLKs *i.e.* BRI1 and FLS2, respectively (Li *et al.*, 2002; Chinchilla *et al.*, 2007), and the same co-receptor BAK1 functions with both of them to provide specificity of response regulation (Wang, 2012; Wang *et al.*, 2014). Additional proteins (*e.g.* BIK1) also form a part of these receptor complexes (Lu *et al.*, 2010). Genetic and functional interactions as well as direct physical associations of plant G-protein complex with RLKs suggest that the G-protein cycle in plants may be activated/regulated by phosphorylation-based mechanisms (Liang *et al.*, 2016; Liang *et al.*, 2018). We have previously shown that plant G-proteins are key regulators of root nodule development in soybean (Roy Choudhury & Pandey, 2013; Roy Choudhury & Pandey, 2015). As the receptors of nodule formation and many of their downstream signaling components are well-established (Endre *et al.*, 2002; Nishimura *et al.*, 2002; Madsen *et al.*, 2003; Downie, 2014; Singh *et al.*, 2014; Ferguson *et al.*, 2018), we chose this system to explore the receptor-dependent regulation of G-protein signaling in plants.

Nodule formation begins with the secretion of flavonoids by roots, which are sensed by compatible rhizobia in soil. Rhizobia release nodulation factors (NFs) which are perceived by the Nod factor receptors (NFRs) present at the plasma membrane of epidermal root hair cells of compatible host plants (Limpens *et al.*, 2003; Madsen *et al.*, 2003). Root hairs deform, entrap rhizobia, which then invade the epidermal cells using infection threads followed by the penetration of root cortical tissue. The colonization of host cells by rhizobia leads to the formation of specialized organelles the ‘symbiosomes’, where rhizobia differentiate into their nitrogen-fixating forms called bacteroids. The multiplication of bacteroids within the dividing cortical space and concomitant organogenesis results in nodule development (Oldroyd & Downie, 2008; *Oldroyd et al., 2011*; Popp & Ott, 2011).

The Nod factor receptors (NFRs) are RLKs, which possess a LysM motif in their extracellular domain (LysM-RLK). In soybean, these are represented by NFR1α and 1β, which are active kinases and NFR5α and 5β, which are the co-receptors of NFR1 and have no kinases activity (Indrasumunar *et al.*, 2010; Indrasumunar *et al.*, 2011). NFRs have been shown to bind NFs and initiate downstream signaling, including changes in calcium levels and transcriptional responses in the nucleus (Desbrosses & Stougaard, 2011; Oldroyd *et al.*, 2011; Popp & Ott, 2011; Broghammer *et al.*, 2012). We have previously shown that the soybean Gα proteins and their regulatory RGS proteins interact with NFR1 (Roy Choudhury & Pandey, 2013; Roy Choudhury & Pandey, 2015). We also showed that NFR1 phosphorylates the RGS proteins, thereby affecting the G-protein cycle indirectly. However, no effect of NFR1 interaction was observed on the GTP-binding or GTPase activity of Gα protein (Roy Choudhury & Pandey, 2015).

To explore the receptor-mediated regulation of G-protein signaling, we focused on additional receptor-like proteins known to be involved in controlling nodulation. The symbiosis receptor kinase (SymRK) is another such protein, which is present at the plasma membrane and acts downstream of NF perception (Endre *et al.*, 2002; Stracke *et al.*, 2002). SymRKs encode a protein with three LRRs in the predicted extracellular region and an intracellular protein kinase domain, with conserved tyrosine-valine residues, typical of many RLKs (Endre *et al.*, 2002; Stracke *et al.*, 2002; Indrasumunar *et al.*, 2015; Perraki *et al.*, 2018). The mechanistic details of SymRKs, their ligands or their direct downstream targets are not known, although many proteins have been shown to interact with them (Kevei *et al.*, 2007; Zhu *et al.*, 2008; Chen *et al.*, 2012; Yuan *et al.*, 2012; Vernie *et al.*, 2016).

In this study, we demonstrate a direct SymRK-mediated regulation of G-protein signaling during nodulation in soybean. We show that the soybean SymRKs (GmSymRKα and GmSymRKβ or NORK) interact with and phosphorylate the Gα protein of the heterotrimeric G-protein complex. Intriguingly, two of the phosphorylated amino acids are in the conserved GTP-binding domain of the Gα protein. Phosphorylation of the serine residues in this site abolishes the ability of Gα to bind GTP, essentially making the protein biochemically inactive. Furthermore, the phospho-mimetic Gα is unable to bind Gβγ proteins, suggesting an altered dynamics and/or availability of the active proteins for downstream signaling. Our results thus uncover a novel G-protein signaling mechanism where not only the activity but also the availability of the signaling proteins is dependent on receptor-mediated phosphorylation of the Gα protein.

## MATERIAL AND METHODS

### Plant material and hairy root transformation

Soybean (*Glycine max*) wild type (‘Williams 82’) seeds were grown on (BM 7 35%) in the greenhouse (16 h light/8 h dark) for 12 days at 25°C. Hairy root transformation was performed as per our previous protocols (Roy Choudhury & Pandey, 2013; Roy Choudhury & Pandey, 2015). Rhizobium strain (USDA136) cultured in Vincent’s rich medium was used for bacterial infection. Nodules were counted at 32 days after rhizobium infection. Three biological replicates were used for each construct. At least 35 to 40 transgenic hairy roots were used in individual experiment for each construct. The data were averaged, and statistically significant values were determined by Dunn’s multiple comparisons test.

### Generation of constructs

The gene fragment for *RNAi* construct of *SymRK* was cloned into the pCR8/GW vector (Invitrogen) and subsequently transferred into binary vector CGT11017A by Gateway-based cloning (Invitrogen). The *RNAi* construct was expressed under the control of the *FMV* promoter with GFP selection marker (Govindarajulu *et al.*, 2009). The mutant versions of soybean *SymRKα* and *Gα* genes were generated by using the Quick-change® Site-Directed Mutagenesis Kit (Agilent). For overexpression, native and mutant versions of *SymRKα*, *Gα* and *RGS* genes were transferred into pCAMGFP-CvMV-GWi vector by Gateway-based cloning (Invitrogen). The overexpression constructs were expressed under the control of the *CvMV* promoter with GFP selection marker (Roy Choudhury & Pandey, 2013). Sequence confirmed RNAi and overexpression constructs including empty vectors (EV), were transformed into *Agrobacterium rhizogenes* strain K599.

### RNA isolation and quantitative Real-time PCR

Total RNA isolation and cDNA synthesis from different root tissues of soybean were performed as described previously (Roy Choudhury *et al.*, 2011). All cDNA samples were diluted 1: 20 in sterile water. qRT-PCR assays were performed with gene-specific primers and soybean *Actin* gene was used as an internal control for transcript levels studies (Bisht *et al.*, 2011). Two biological replicates were used for gene expressions analysis. The qRT-PCR was repeated three times for each biological replicate and data were averaged.

### Protein-protein interaction

The interaction between SymRKα and SymRKβ with Gα (Gα1-4) was performed using the mating-based yeast split-ubiquitin system. Briefly, SymRKα and SymRKβ (full length, N- or C-terminal regions) were fused with the C-terminal half of ubiquitin (CUb fusions). Gα were fused with the N-terminal half of ubiquitin (NUb fusions). NUbwt fusion constructs, which show intrinsic interaction with CUb fusion constructs, were used as positive controls and empty vector (EV) was used as negative control. Yeast transformations and mating were performed as described previously (Pandey & Assmann, 2004). To study the interaction between native and mutant Gα with native RGS or Gβ, Gα genes were used as NUb fusions and RGS or Gβ was used as CUb fusions. All experiments were repeated at least two times and a representative image is shown.

For bimolecular fluorescence complementation (BiFC) assays, the *Gα1-4* genes were cloned into 77 nEYFP-N1 vector (containing EYFP at the C-terminal end), and the *SymRKα* genes, *RGS2* gene or *Gβ* genes were cloned into 78cEYFP-N1vector (containing cEYFP at the C-terminal end). For tri-partite interaction between Gα, Gβ and Gγ, specific *Gγ* genes (*Gγ1, Gγ5 and Gγ9*) were cloned in pEarlyGate203 vector. All constructs were transformed into *Agrobacterium tumefaciens* strain GV3101. Abaxial surface of tobacco leaves was infiltrated with *A. tumefaciens* containing either the gene of interest in different combinations or EV as negative control. Infiltrated plants were incubated in darkness for 24 h followed by 48 h incubation in light. The leaves were imaged with the Nikon Eclipse E800 microscope with epi-fluorescence module for YFP fluorescence detection as previously reported (Roy Choudhury *et al.*, 2012). Experiments were repeated two times and each time at least three independent infiltrations were performed for each construct.

For co-immuno precipitation (co-IP) of proteins, specific proteins were expressed as epitope tagged versions. To generate 35S:Flag-Gα1 and 35S:Myc-SymRKα constructs, the full-length cDNAs of soybean Gα1 and SymRKα were obtained by PCR amplification and subcloned into pEarlygate 202, and pEarlygate 203 vectors, respectively. Selected constructs were introduced into *Agrobacterium tumefaciens* strain GV3101 and the bacterial suspensions were infiltrated into the abaxial surface of 4-week old tobacco (*Nicotiana tabacum*) leaves. After 96 h, proteins were extracted from inoculated leaves in the extraction buffer as described previously (Roy Choudhury & Pandey, 2015). The homogenate was centrifuged at 8000 *g* for 10 min to remove debris. For each immunoprecipitation experiment, ~200 μg of protein extracts were incubated with anti-Myc antibody in the presence of binding buffer as described previously (Roy Choudhury & Pandey, 2015). Finally, pulled-down proteins were separated on SDS-PAGE and analyzed by immunoblotting with anti-Flag and anti-Myc antibodies (Sigma-Aldrich).

### Recombinant protein purification

Native and mutant versions of full-length Gα1 and C-terminal region of RGS2 and SymRKα proteins were cloned into the pET-28a vector (Novagen, Gibbstown, NJ, USA) and recombinant proteins were purified using Ni^2+^-affinity chromatography as previously described (Roy Choudhury *et al.*, 2013). Protein aliquots were snap-frozen in liquid nitrogen and stored at −80°C for *in vitro* phosphorylation assay and GTPase activity assay.

### *In vitro* phosphorylation assay and LC-MS/MS analysis

For *in vitro* phosphorylation assays, 1.8 μg kinase (C-terminal region of SymRKα or NFR1α) and 1 μg substrate proteins (native or mutant Gα) were incubated in 20 μl of 25 mM TrisCl (pH 8.0), 15 mM MgCl_2_, 2 mM MnCl_2_, 7.5 mM NaCl, 0.05 mM EDTA, 0.05 mM DTT, 5% glycerol and 75 μM ATP at 27°C for 30 min. The reaction was stopped by adding 5X SDS sample loading buffer followed by boiling for 5 min. Samples were analyzed via 10% SDS-PAGE. The gel was stained with Pro-Q® Diamond phosphoprotein gel stain (Invitrogen) and imaged using the Typhoon 9410 variable mode imager (Molecular Dynamics). For protein staining, this same gel was immersed in SYPRO™ ruby protein gel stain (Invitrogen) followed by imaging using the Typhoon 9410 variable mode imager.

For LC-MS/MS analysis, 20 μL of the kinase-substrate protein solution was reduced (10 mM TCEP) and alkylated (20 mM Iodoacetemide) before digestion with trypsin (0.2 μg) overnight. Digest was acidified with 1% TFA before clean up with C18 tip. The extracted peptides were dried and each sample was resuspended in 20 μL 5% ACN/ 0.1% FA. Two μL (~1μg) peptides were analyzed by LC-MS with a Dionex RSLCnano HPLC coupled to a LTQ-Orbitrap Velos Pro (Thermo Scientific) mass spectrometer using a 2 h gradient. Peptides were resolved using 75 μm × 25 cm PepMap C18 column (Thermo Scientific). All MS/MS samples were analyzed using Mascot (Matrix Science, London, UK; version 2.5.1.0). Mascot was set up to search the provided sequence as well as common contaminants. The digestion enzyme was set as trypsin. Mascot was searched with a fragment ion mass tolerance of 0.60 Da and a parent ion tolerance of 10 ppm. Oxidation of methionine, carbamidomethylation of cysteine, acetylation of N-terminal of protein and phosphorylation of serine, threonine and tyrosine were specified in Mascot as variable modifications. Scaffold (version Scaffold_4.5.3 Proteome Software Inc., Portland, OR) was used to validate MS/MS based peptide and protein identifications. Peptide identifications were accepted if they could be established at greater than 80.0% probability by the Peptide Prophet algorithm (Keller *et al.*, 2002) with Scaffold delta-mass correction. Protein identifications were accepted if they could be established at greater than 99.0% probability and contained at least 2 identified peptides. Protein probabilities were assigned by the Protein Prophet algorithm (Nesvizhskii *et al.*, 2003). Proteins that contained similar peptides and could not be differentiated based on MS/MS analysis alone were grouped to satisfy the principles of parsimony. Proteins sharing significant peptide evidence were grouped into clusters.

### GTP-binding and GTPase activity assay

Real-time fluorescence-based GTP-binding and GTP-hydrolysis assays of native and mutant Gα proteins were performed using BODIPY-GTP FL (Invitrogen) as described previously (Roy Choudhury *et al.*, 2013). Assays were performed at 25°C in a 200 μl reaction volume in assay buffer (20 mM Tris, pH 8.0 and 10 mM MgCl_2_). GTPase activity of native and mutant Gα proteins were detected with or without an RGS protein using the ENZchek phosphate assay kit (Invitrogen) as previously reported (Roy Choudhury *et al.*, 2013). All assays were measured by Infinite M200 Pro (Tecan).

## RESULTS

### Soybean Gα proteins interact with symbiosis receptor kinase (SymRK)

Heterotrimeric G-protein regulate nodule formation (De Los Santos-Briones *et al.*, 2009; Roy Choudhury & Pandey, 2013; Roy Choudhury & Pandey, 2015). As per the established paradigm of G-protein signaling, the Gα proteins are expected to function with the receptor-like proteins localized at the plasma membrane. In our previous experiments, even though we saw an interaction between the NFR1 and Gα proteins, a direct effect of this interaction on Gα activity was not observed. As Gα proteins in plants are known to interact with a variety of RLKs (Llorente *et al.*, 2005; *Liu et al., 2013*; Aranda-Sicilia *et al.*, 2015; Tunc-Ozdemir *et al.*, 2016; Zhou *et al.*, 2019), we evaluated interactions between additional plasma membrane-localized, receptor-like proteins that are known to be involved in plant-microbe interaction (*e.g.* NFR1α and β, NFR5 α and β, SymRK α and β, NARK etc.) with different G-protein components. In our initial screen, SymRKs, which are central to nodulation signaling pathways, interacted with the Gα proteins. SymRK is proposed to work downstream of the NFR1 proteins, although their ligands or direct downstream targets are not well established (Endre *et al.*, 2002; Stracke *et al.*, 2002; Holsters, 2008; Markmann *et al.*, 2008). The soybean genome encodes two SymRK homologs, SymRKα and SymRKβ, which are 95% identical. SymRKs have an extracellular malectin-like domain (MLD), a region with leucine-rich repeats (LRR), and an intracellular region containing a conserved Ser/Thr protein kinase domain, connected with one transmembrane region (Indrasumunar *et al.*, 2015; Li, H *et al.*, 2018). Both genes show similar transcript levels in different tissue types (Fig. S1). To corroborate our preliminary observation, we evaluated the full-length, C-terminal and N-terminal regions of SymRKα and the C-terminal region of SymRKβ for their interaction with the four soybean Gα proteins in a split-ubiquitin-based interaction assay. For these interactions, the SymRK proteins were expressed as bait proteins (CUb fusions) and the full-length Gα proteins (Gα1, Gα2, Gα3 and Gα4) were expressed as prey proteins (NUb fusions) along with appropriate positive and negative controls. The Gα proteins interacted with full-length as well as C-terminal region of SymRKs, but not with the N-terminal region as assessed by yeast growth on media lacking Leu, Trp, His and Ade, in the presence of 250 μM Met (Fig. 1a, S2a). BiFC assays were performed to test *in planta* interaction between these proteins. For these assays, the Gα proteins were expressed as a fusion protein with nEYFP at its C-terminal end and SymRKα was expressed as a fusion protein with cEYFP at its C-terminal end. Strong fluorescence was detected when both constructs were co-expressed in tobacco leaves, confirming that the Gα proteins interact with SymRK proteins (Fig. 1b). To validate the interaction specificity, we also tested the interactions of Gα1 with NFR1α (positive control), NFR5α (negative control), N-terminal region of RGS2 protein (negative control) or with an empty vector (EV, negative control). A strong signal was observed when Gα was co-expressed with NFR1α protein but not when EV control, NFR5α or N-terminal region of RGS2 was used as interaction partners (Fig. 1b). Gα1 also interacted strongly with the C-terminal of SymRKβ (Fig. S2b). To corroborate these interactions further, we performed in vivo co-IP assays with Flag epitope-tagged Gα1 and Myc epitope-tagged SymRKα proteins expressed in tobacco leaves. Anti-Myc antibodies could pull down Flag-tagged Gα1 proteins in these assays confirming the in planta interaction between them (Fig. 1c). These data establish that the soybean Gα proteins interact with the SymRK proteins.

**Fig. 1.**
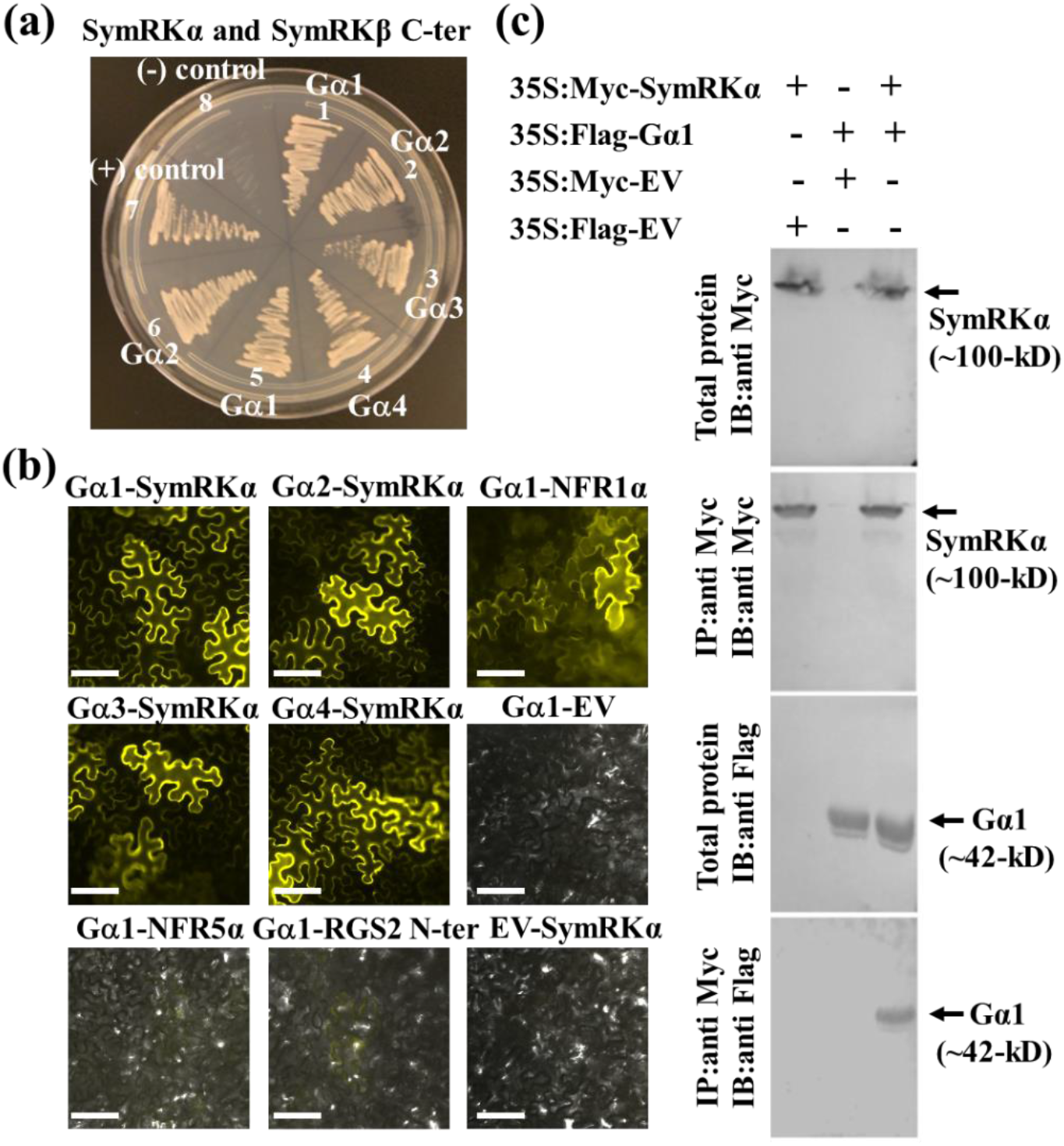
Gα proteins interact with SymRK. (a) Interaction between Gα proteins (1-4) with the C-terminal regions of SymRKα and SymRKβ proteins using split-ubiquitin based interaction assay. The picture shows yeast growth on selective media with 250 μM methionine. In all cases, Gα proteins were used as NUb fusions and C-terminal SymRK proteins were used as CUb fusions. Three biological replicates of the experiment were performed with identical results. The combinations were (1) SymRKα-CUb+Gα1-Nub, (2) SymRKα-CUb+Gα2-Nub, (3) SymRKα-CUb+Gα3-Nub, (4) SymRKα-CUb+Gα4-Nub, (5) SymRKβ-CUb+Gα1-Nub, (6) SymRKβ-CUb+ Gα2-Nub, (7) SymRKα-CUb+ Gα1-NUbwt (positive control), (8) C-terminal SymRKα-CUb+ NUb-vector (negative control). (b) Interaction between Gα proteins (Gα1, Gα2, Gα3, Gα4 in 77-nEYFP-N1) with SymRKα, NFR1α, NFR5α, RGS2 N-terminal or EV (in 78-cEYFP-N1) proteins using bimolecular fluorescence complementation (BiFC) assay. Agrobacteria containing different combinations were infiltrated in tobacco leaves. The reconstitution of YFP fluorescence due to protein-protein interaction was visualized. Interaction between Gα and NFR1α was used as a positive control and interaction between Gα and NFR5α Gα and N-terminal RGS2, Gα+ EV and SymRKα +EV were used as negative controls. At least five independent infiltrations were performed for each protein combination with similar results. Bar = 50 μm. (c) Interaction between Gα1 and SymRKα protein using an in vivo co-IP assay. Anti-Myc antibody was used to pull down Flag-tagged Gα1 from total protein extracts of plants expressing 35S:Flag-Gα1 and 35S:Myc-SymRKα (lane 3) but not from total protein extracts from plants expressing 35S:Flag-Gα1 and 35S:Myc-tagged EV (lane 2) or 35S:Myc-SymRKα and 35S:Flag-tagged EV (lane 1).

### SymRK phosphorylates Gα proteins

The interaction between Gα and SymRK proteins, SymRKs being active protein kinases (Saha *et al.*, 2016; Vernie *et al.*, 2016), and the possibility that Gα proteins are potential substrates for RLKs in plants (Liu *et al.*, 2013; Tunc-Ozdemir *et al.*, 2016; Chakravorty & Assmann, 2018; Li, B *et al.*, 2018; Zhou *et al.*, 2019), prompted us to evaluate if SymRKs can directly phosphorylate Gα proteins. We performed *in vitro* phosphorylation assays using recombinant, full-length Gα1 protein as a substrate and C-terminal regions of SymRKα and NFR1α proteins as kinases. As we have reported previously, (Roy Choudhury & Pandey, 2015) NFR1α was not able to phosphorylate Gα protein however, strong phosphorylation of Gα was observed when SymRKα was used as a kinase (Fig. 2a). Both NFR1α and SymRKα were also autophosphorylated under these assay conditions (Fig. 2a).

**Fig. 2.**
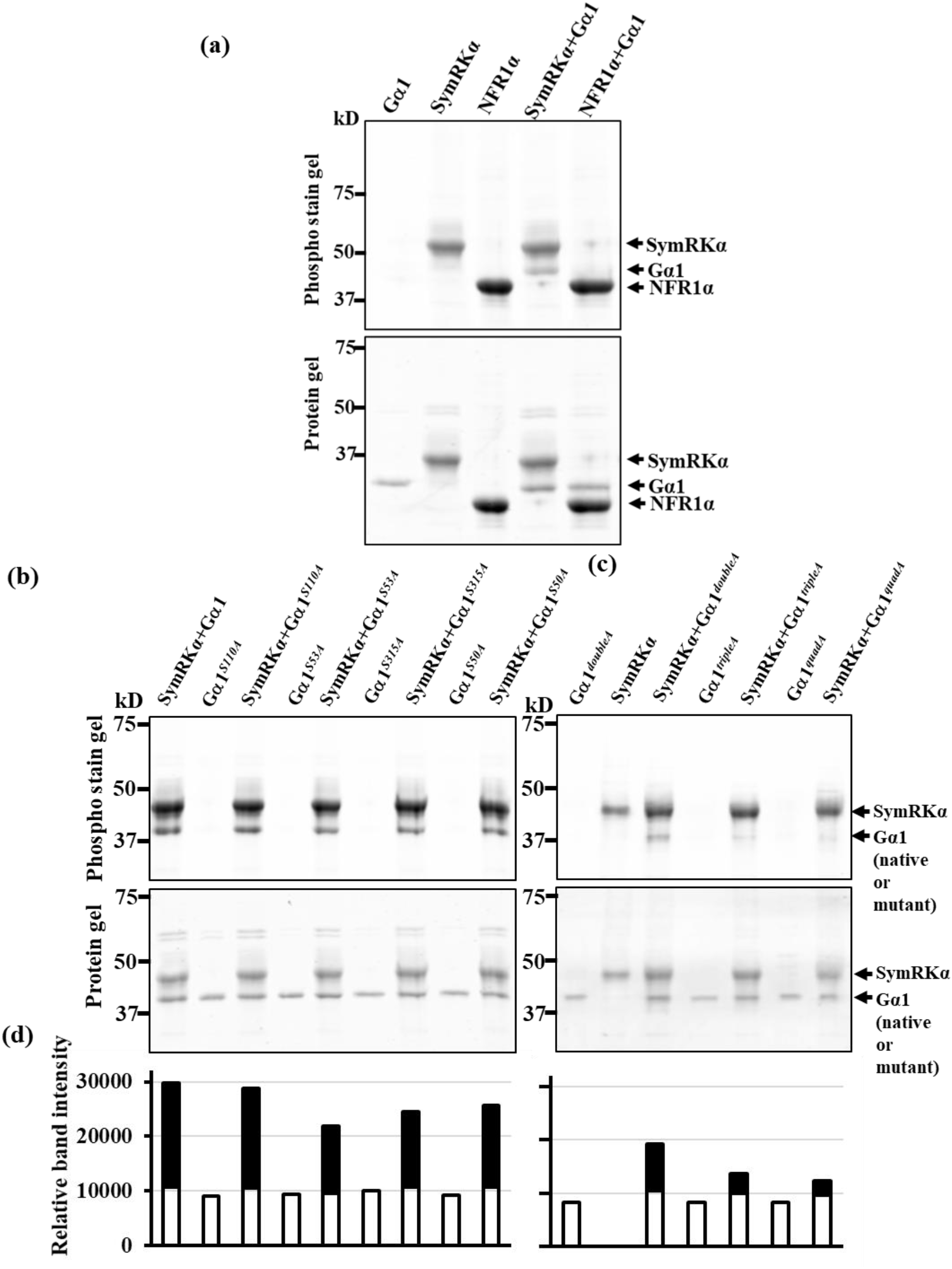
Active SymRKα phosphorylates Gα protein. (a) Gα protein phosphorylation by SymRKα. Gα1 protein was incubated with SymRKα C-terminal or NFR1α C-terminal protein for *in vitro* phosphorylation assay. (b) Phosphorylation assay using single mutations in Gα1 protein (based on the information from LC-MS data). (c) Phosphorylation assay using higher order mutations in Gα1 protein. In (b) and (c), recombinant purified proteins (single mutants: Gα1^*S50A*^, Gα1^*S53A*^, Gα1^*S110A*^, Gα1^*S315A*^); higher order mutants: Gα1^*doubleA*^ (Gα1^*S53, S110A*^), Gα1^*tripleA*^ (Gα1 ^*S53A, S110A, S315A*^) and Gα1^*quadA*^ (Gα1^*S50A, S53A S110A, S315A*^) were incubated with SymRKα C-terminal protein for *in vitro* phosphorylation. In (a), (b) and (c), the upper panel represents Pro-Q® Diamond phospho-stained gel and the lower panel shows the same gel stained with Sypro ruby to visualize protein profiles. (d) The quantification of band intensity (b) and (c) of phosphorylated Gα1 (upper panel) and Gα1 protein (lower panel) by Image J.

LC-MS/MS analysis identified four phosphorylation sites (S50, S53, S110 and S315) in the Gα1 protein, which were phosphorylated by SymRKα, at a minimum localization threshold of 99% (Fig. S3). These sites are conserved in all four Gα proteins of soybean (Bisht *et al.*, 2011; Roy Choudhury *et al.*, 2014). To validate the LC-MS/MS data, we generated point mutations at each of these residues by changing the serine (S) to alanine (A) in Gα1 (phospho-dead version) and evaluated them for their ability to be phosphorylated by SymRKα. Phosphorylation of Gα was marginally reduced when any of the single point-mutant variants of the protein were used, suggesting that the protein is phosphorylated at multiple sites (Fig. 2b, d). We also generated phospho-dead versions of Gα1 protein at additional sites in conjunction with Gα1^*S53A*^ i.e. double (Gα1^*S53A, S110A*^), triple (Gα1^*S53A*, *S110, S315A*^) and quadruple (Gα1^*S53A*, *S110A, S315A*,*S50A*^ or Gα1^*quadA*^) and tested the recombinant proteins in phosphorylation assays. In comparison to any single Gα1 variant, each additional mutation lead to significantly reduced phosphorylation with negligible phosphorylation observed in Gα1^*quadA*^, suggesting that under *in vitro* conditions, each of these four serine residues of Gα proteins are phosphorylation targets of SymRK (Fig. 2c, d).

### Phosphorylated Gα protein cannot bind GTP

Intriguingly, two of the phospho-sites in Gα protein S50 and S53 mapped to its GTP-binding region, which is critical for its activity (Bisht *et al.*, 2011; Roy Choudhury *et al.*, 2014). To evaluate the effects of phosphorylations on the GTP-binding activity, we generated the phospho-mimic versions of Gα1 protein by replacing the serine residues with aspartic acid (D) at each of the potential phosphorylation sites (Gα1^*S50D*^, Gα1^*S53D*^, Gα1^*S110D*^ and Gα1^*S315D*^). We also generated a protein with all four serine phospho-sites converted to aspartic acid (Gα1^*S110D*, *S53D*, *S315D*, *S50D*^ or Gα1^*quadD*^). Native and phospho-mimic Gα1 recombinant proteins were evaluated for GTP-binding and -hydrolysis using real-time BODIPY fluorescence-based assays (Bisht *et al.*, 2011). A constitutively active version of Gα protein, Gα1^*Q223L*^, which exhibits GTP-binding but no hydrolysis (Roy Choudhury *et al.*, 2014; Roy Choudhury & Pandey, 2017) was used as a control. Native Gα1 showed the expected GTP-binding and -hydrolysis, whereas only GTP-binding was detected in Gα1^*Q223L*^. The variant proteins Gα1^*S110D*^ and Gα1^*S315D*^ exhibited GTP-binding and hydrolysis similar to the native Gα1, however the variants Gα1^*S50D*^, Gα1^*S53D*^ and Gα1^*quadD*^ exhibited no GTP-binding (Fig. 3a).

**Fig. 3.**
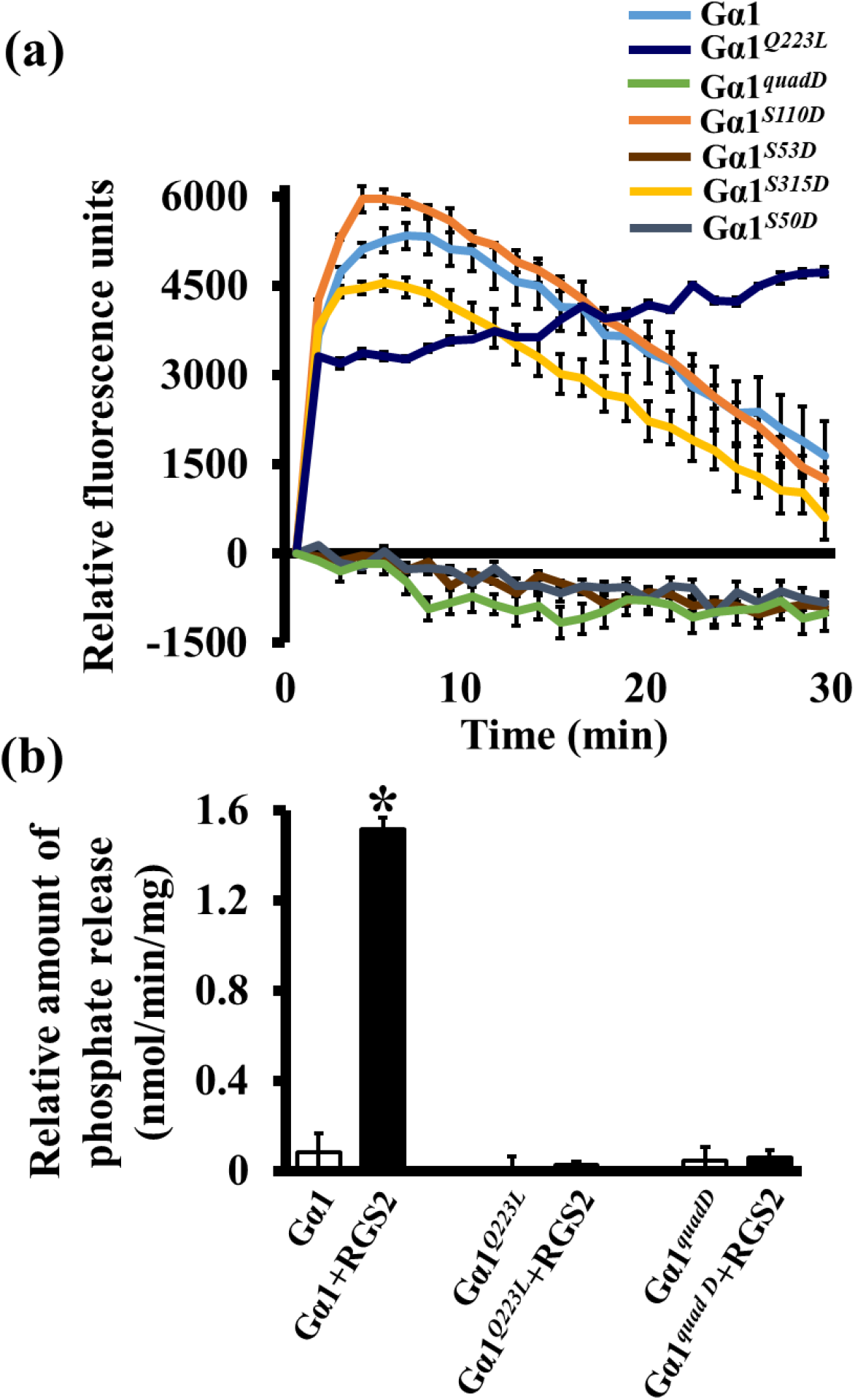
Changes of GTP-binding and GTPase activity of phospho-mimetic Gα. (a) GTP-binding and GTP-hydrolysis of native and mutant Gα1 proteins were detected by GTP-BODIPY-FL in a real time fluorescence based assays. GTP-binding and GTP-hydrolysis of native Gα1, constitutively active Gα1 (Gα1^*Q222L*^) and phospho-mimetic Gα1 proteins (Gα1^*S50D*^, Gα1^*S53D*^, Gα1^*S110D*^, Gα1^*S315D*^ and Gα1^*quadD*^) were measured. The upward slope represents GTP-binding whereas the downward slope represents GTP-hydrolysis. Data are one of two independent experiments, each with three replicates, mean ± S.E. (b) Rate of Pi release due to the GTPase activity of native, constitutively active (Gα1^*Q223L*^) and phospho-mimetic Gα1 (Gα1^*quadD*^) in the absence or presence of native RGS2 protein. Experiments were repeated three times, and data were averaged. Error bars represent the mean ±S.E.

To examine the GTPase activity of the native and variant Gα proteins, we performed a phosphate (Pi) release assay in the absence or presence of an RGS protein (Roy Choudhury *et al.*, 2012). Native Gα1 protein as well as the variants Gα1^*S110D*^ and Gα1^*S315D*^ exhibited slow phosphate release, which significantly increased in the presence of RGS2 protein (Fig. 3b, S4). The variant Gα1^*Q223L*^ showed no Pi release either by itself or in the presence of RGS protein (Fig. 3b). Similarly, no Pi release was observed in the protein variants Gα1^*S50D*^, Gα1^*S53D*^ and Gα1^*quadD*^ (Fig. 3b, S4). Overall, these data establish that SymRKs phosphorylate Gα proteins in their active site, which causes a complete loss of their GTP-binding activity.

### Overexpression of phospho-mimetic *Gα* increases nodule number in plants

To determine the effects of Gα phosphorylation on nodule formation *in planta*, the phospho-deficient and phospho-mimetic versions of Gα1 along with native Gα1 and RGS2 protein (as control) were overexpressed in soybean hairy roots (Roy Choudhury & Pandey, 2013; Roy Choudhury & Pandey, 2015). Transcript levels of corresponding genes were tested to ascertain their higher expression (Fig. S5a, b). We first evaluated the number of deformed root hairs and nodule primordia in overexpression roots at 4 dpi and 6 dpi after *B. japonicum* infection, respectively. Both deformed root hairs and nodule primordia numbers were significantly reduced due to the overexpression of native *Gα1* or *Gα1*^*quadA*^ as compared to the *EV* control hairy roots. Interestingly, opposite trends were seen with the overexpression of *Gα1*^*quadD*^ as these hairy roots had higher numbers of deformed root hairs and nodule primordia compared with the *EV* hairy roots; which was also the case with the roots overexpressing *RGS2* (used as a positive control). Compared to the *EV* hairy roots ~35% fewer deformed root hairs and nodule primordia were observed in *Gα1* and *Gα1*^*quadA*^ overexpressed hairy roots; whereas ~50% more deformed root hairs and ~55% more nodule primordia were detected in *Gα1*^*quadD*^ and *RGS2* overexpression hairy roots, respectively (Fig. 4a, b).

**Fig. 4.**
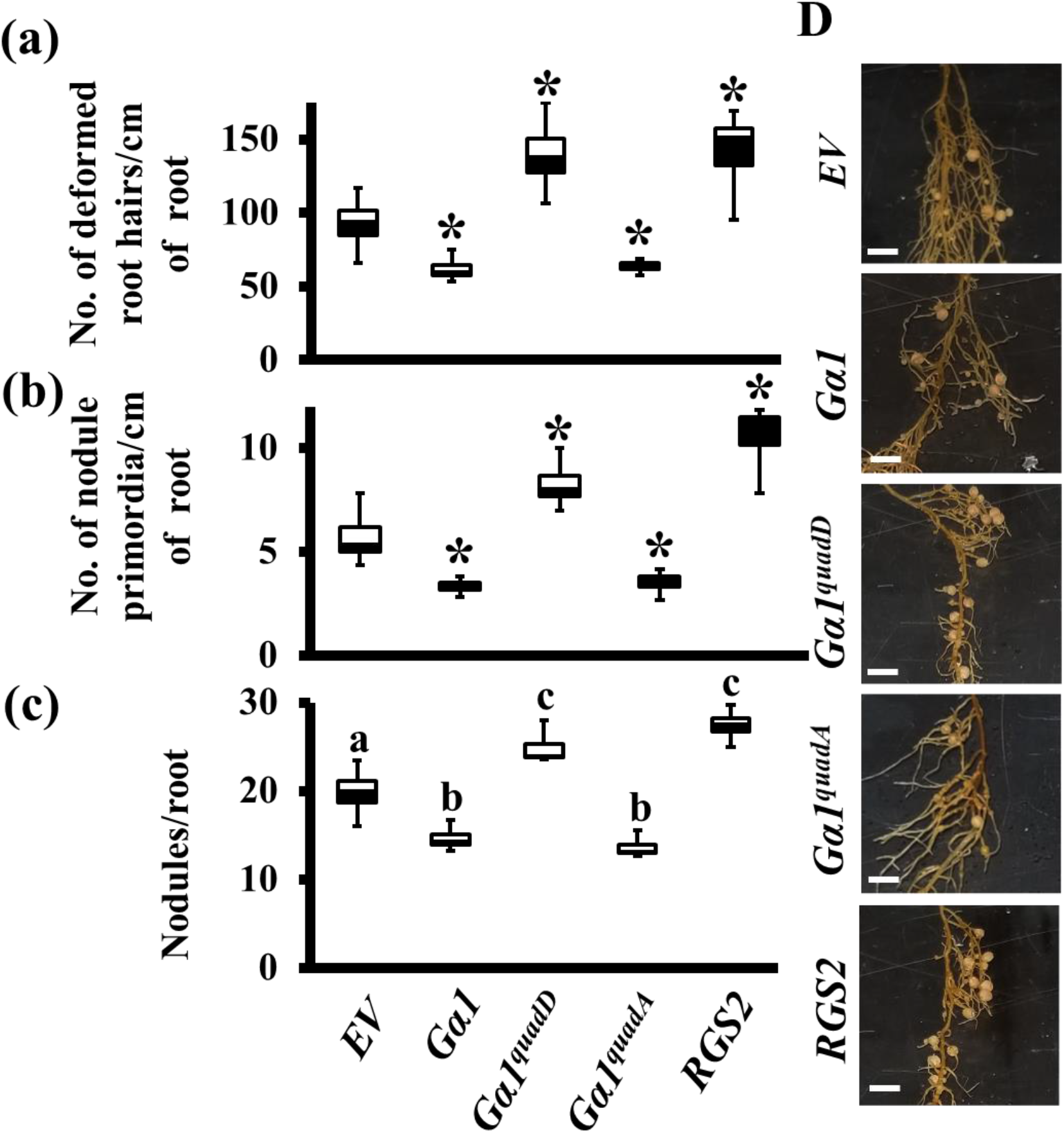
The effect of overexpression of phospho-dead and phospho-mimetic versions of Gα on nodulation. (a) Native, *Gα1*^*quadD*^ and *Gα1*^*quadA*^ versions of *Gα1* and native *RGS2* genes driven by *CvMV* promoter were used for hairy root transformation. Quantification of deformed root hairs/cm in overexpression roots compared with empty vector (*EV*) at 4 dpi with *B. japonicum*. (b) Quantification of nodule primordia/cm in over-expression roots compared with *EV* at 6 dpi with *B. japonicum*. The data presented in (a) and (b) are average values from three independent experiments (n =10). Error bars represent ±S.E. Asterisks (*) indicate a significant differences (* = P < 0.05; Student’s *t* test). (c) Quantification of nodule number in over-expression roots compared with *EV* at 32 dpi with *B. japonicum*. The data represent average of 3 biological replicates (35-40 individual plants/biological replicate) containing transgenic nodulated roots. Different letters indicate significant differences (Dunn’s multiple comparisons test, P < 0.05) between samples. (d) Representative pictures of transgenic hairy roots overexpressing native, *Gα1*^*quadD*^, *Gα1*^*quadA*^ versions of *Gα1* and native *RGS2* compared to *EV* control lines at 32 dpi with *B. japonicum*. Bar = 50 mm.

We also evaluated the nodule numbers in these hairy roots. As we have reported previously, the number of nodules were significantly reduced and increased due to the overexpression of native *Gα1* and *RGS2*, respectively (Roy Choudhury & Pandey, 2015). Overexpression of *Gα1*^*quadA*^ showed similar results as the native Gα, with ~30% reduction in nodule number compared to the *EV* containing hairy-roots. In contrast, the overexpression of *Gα1*^*quadD*^ resulted in the opposite phenotypes with more nodules (~25%) formed per root compared to the *EV* hairy roots (Fig. 4c, d). These data suggest that SymRK-mediated phosphorylation of Gα protein is an important regulatory mechanism during nodule formation in soybean as the overexpression of a phospho-mimic Gα results in phenotypes opposite to the overexpression of a native Gα.

We further assessed the effect of phosphorylated Gα proteins on nodule formation in the context of SymRK-dependent transcriptional regulation. We first corroborated the role of SymRK as an important positive regulator of nodulation by altering its expression or activity in transgenic hairy roots. We generated *SymRK-RNAi* roots (*sym*) in which both *SymRKα* and *SymRKβ* transcripts were significantly downregulated (Fig. S6a). In addition, we generated transgenic hairy roots overexpressing native *SymRKα* or a point mutant *SymRKα^K617E^*, which has no kinase activity (Fig. S6b). Expression of both native and mutant *SymRK* genes was higher than in *EV*-containing hairy roots (Fig. S6c). Quantification of nodule numbers confirmed that SymRKs are positive regulators of nodule formation as the roots expressing lower (*RNAi*) or higher (overexpression) levels of the gene had significantly fewer or more nodules, respectively (Fig. 5a, b, c). The kinase activity of SymRK was essential for this effect as the overexpression of *SymRKα*^*K617E*^ did not result in the production of more nodules (Fig. 5b, c).

**Fig. 5.**
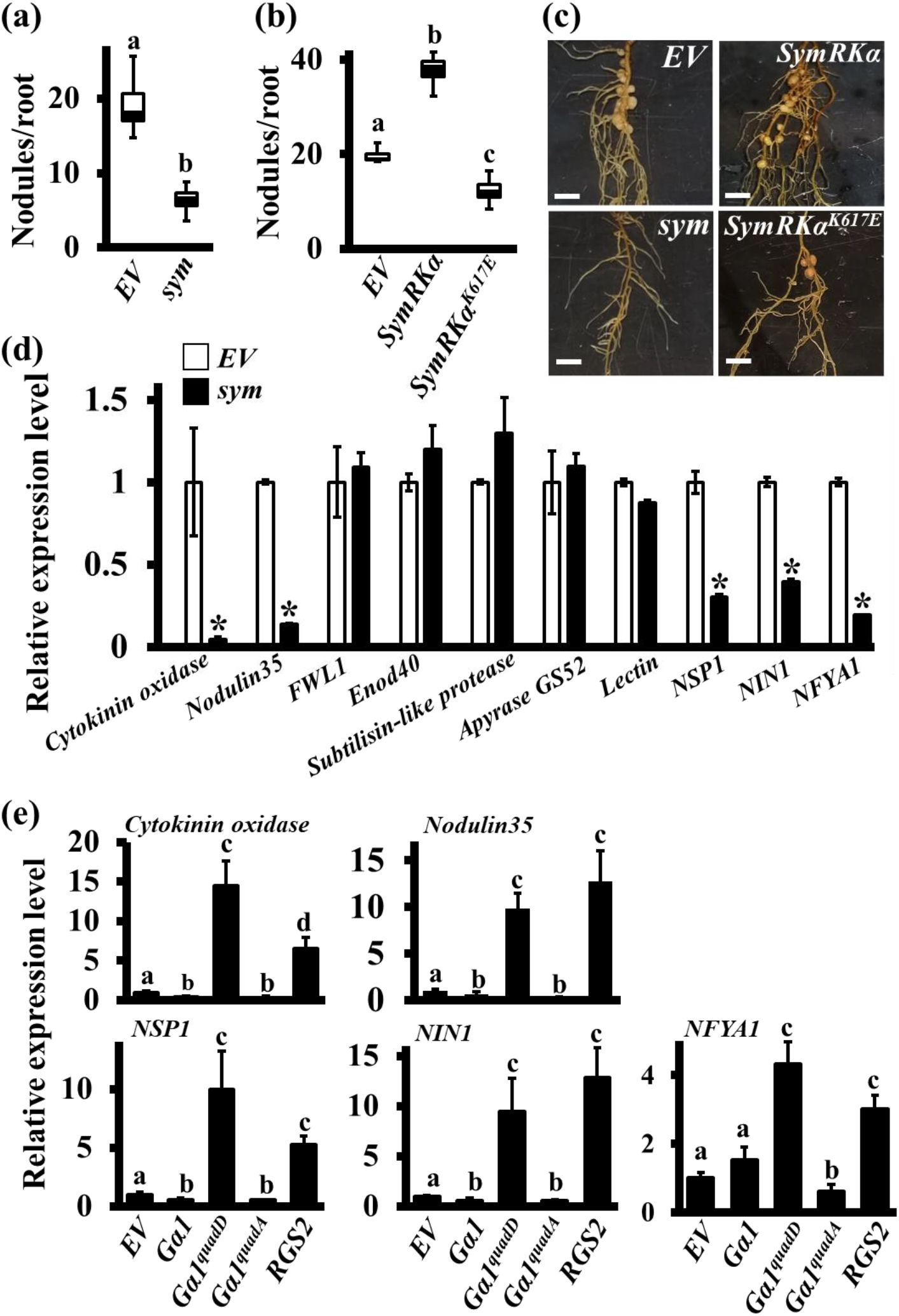
Nodulation phenotypes of *sym* mutants and expression of nodulation-related genes in *SymRK-RNAi* roots and in roots overexpressing native *Gα1*, *Gα1*^*quadD*^ and *Gα1*^*quadA*^. (a) Nodulation phenotype of *SymRK* knockdown hairy roots and (b) hairy roots overexpressing native and mutant *SymRKα* genes. Nodule numbers on transgenic hairy roots were counted at 32 dpi with *B. japonicum* and were compared with their respective *EV* transformed control hairy roots. The data represented in (a) and (b) are average values from 3 biological replicates (35-40 individual plants/biological replicate). Different letters indicate significant differences (Dunn’s multiple comparisons test, P < 0.05) between samples. (c) Representative pictures of *SymRK* knockdown, native *SymRKα* overexpressing lines and mutant *SymRK*^*αK617E*^ overexpressing lines transgenic hairy roots compared to *EV* control roots at 32 dpi with *B. japonicum*. Bar = 50 mm. (d) Relative expression of nodulation-related genes in *EV* (control) and *SymRK-RNAi* hairy roots at 4 dpi after inoculation with *B. japonicum*. Gene specific primers were used to amplify and quantify the transcript levels of *Cytokinin oxidase (Glyma17g054500.1)*, *Nodulin35 (Glyma10g23790.1)*, *FWL1 (Glyma09g187000)*, *Subtilisin-like protease (Glyma17G133400*), *Enod40 (Glyma01g03470.1)*, *Apyrase GS52 (Glyma16g04750), Lectin (Glyma10g113600), NSP1 (Glyma07g04430)*, *NIN1 (Glyma14g00470)* and *NFYA1 (Glyma10g10240).* (e) Relative expression of nodulation-related genes in *EV* (control), *Gα1*, *Gα1*^*quadD*^ and *Gα1*^*quadA*^ and *RGS2* overexpression roots at 4 dpi after inoculation with *B. japonicum*. The expression values across different samples are normalized against soybean *Actin* gene expression. Expression in *EV* roots was set at 1. Error bars represent the ±SE of the mean. Different letters indicate significant differences compared between them (*P < 0.05) using Student’s *t* test.

We tested the expression of ten known nodulation-related marker genes in the hairy roots of *sym* plants. Five of these genes, a *cytokinin oxidase*, *nodulin 35*, *NSP1*, *NIN1* and *NFYA1* exhibited a SymRK-dependent decrease in expression levels, whereas no significant differences were seen in the other five genes (Fig. 5d). We next evaluated the expression of these five genes in transgenic hairy roots overexpressing native, phospho-dead (Gα1^*quadA*^) or phospho-mimetic (Gα1^*quadD*^) versions of Gα1, along with the plants expressing an empty vector or an RGS2 protein as negative and positive controls, respectively. All five genes showed higher expression in *Gα1*^*quadD*^ and *RGS2* overexpressing roots and lower or unaltered expression in roots overexpressing native *Gα1* or *Gα1*^*quadA*^ protein (Fig. 5e), implying that SymRK and Gα proteins share similar gene regulatory pathways during control of nodule formation.

### Phosphorylated Gα proteins are unable to interact with the Gβγ dimer

The inability of the phosphorylated Gα proteins to bind GTP (and therefore exhibiting any canonical G-protein activity), but maintaining the ability to affect the plant phenotypes (*i.e.* increased nodule formation in plants overexpressing Gα^*quadD*^), suggests that SymRK-mediated phosphorylation of Gα protein is biologically important. Two interrelated aspects of Gα proteins define its signaling cycle, its biochemical activity and its ability to interact with the Gβγ dimer or the regulatory RGS protein. Since the phosphorylated protein lacks any biochemical activity, the effect cannot be due to the alterations in the rate or regulation of the G-protein signaling cycle per se. We therefore tested whether the interaction of Gα with its receptors, RGS or Gβγ proteins were altered due to its phosphorylation. Specifically, we evaluated the interactions of Gα1, Gα1^*quadD*^ and Gα1^*quadA*^ proteins with Gβ, RGS2, NFR1α and SymRKα proteins.

To test interaction between Gα and Gβ, the full-length proteins were expressed as prey (NUb fusions) and bait (CUb fusions), respectively, in the split-ubiquitin interaction system. In this assay, all native Gβ proteins (Gβ1-4) interacted with the native Gα1, and with the phospho-dead Gα1 (Gα1^*quadA*^) but intriguingly, no interaction was observed with Gα1^*quadD*^ (Fig. 6a). To corroborate these interactions *in planta*, native Gα1, Gα1^*quadA*^ or Gα1^*quadD*^ proteins were transiently co-expressed with Gβ and Gγ proteins in tobacco leaves. We used two soybean Gβ proteins (Gβ2 and Gβ3), which belong to two different subgroups and three different Gγ proteins (Gγ1, Gγ5 and Gγ9), representing each of the Gγ subgroups (Roy Choudhury *et al.*, 2011) to test the tripartite interaction of Gβγ with Gα1, Gα1^*quadA*^ or Gα1^*quadD*^. In these assays too, Gα1^*quadD*^ did not show any interaction, while the native Gα1 as well as Gα1^*quadA*^ exhibited strong interactions with all tested combinations of Gβγ proteins (Fig. 6b, S7).

**Fig. 6.**
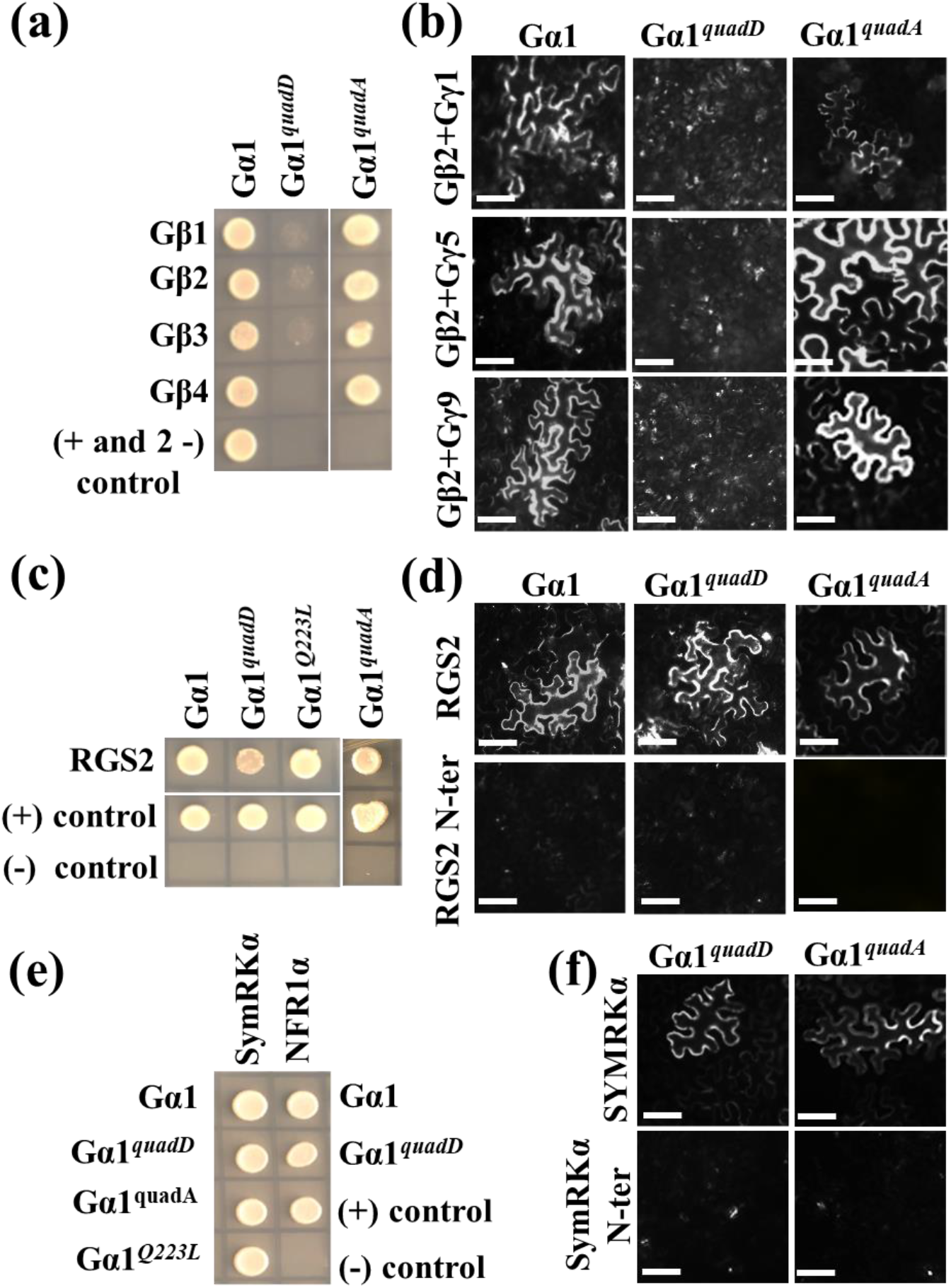
phospho-mimetic Gα does not interact with Gβ. (a) Interaction between Gα1, *Gα1*^*quadD*^ and *Gα1*^*quadA*^ with Gβ (1-4) proteins using split-ubiquitin based interaction assay. In all cases, Gα1 proteins were used as NUb fusions and Gβ proteins were used as CUb fusions. (b) Interaction between Gα1, Gα1^*quadD*^ and Gα1^*quadA*^ (in 77-nEYFP-N1) with Gβ2 (in 78-cEYFPN1) and Gγ (1 or 5 or 9) proteins using BiFC assay. Agrobacteria containing different combinations of Gα1, Gβ and Gγ were infiltrated in tobacco leaves and reconstitution of YFP fluorescence due to protein-protein interaction was visualized. (c) Interaction between Gα1, Gα1^*quadD*^, Gα1^*Q223L*^ and Gα1^*quadA*^ with native RGS2 using split-ubiquitin based interaction assay. In all cases, Gα1 proteins were used as NUb fusions and RGS2 was used as CUb fusions. (d) Interaction between Gα1, Gα1^*quadD*^ and Gα1^*quadA*^ (in 77-nEYFP-N1) with full length and N-terminal RGS2 (in 78-cEYFP-N1) proteins using BiFC assay. Agrobacteria containing different combinations of Gα1 and RGS2 were infiltrated in tobacco leaves and reconstitution of YFP fluorescence due to protein-protein interaction was visualized. (e) Interaction between Gα1, Gα1^*quadD*^, Gα1^*quadA*^ and Gα1^*Q223L*^ with full-length SymRKα and NFR1α. In all cases, Gα1s were used as NUb fusions and full-length SymRKα or NFR1α were used as CUb fusions. In all yeast based interactions (a, c and e), NUbwt fusion and NUb-vector fusion constructs were used as positive and negative controls, respectively. The picture shows yeast growth on selective media with 250 μM methionine. Two biological replicates of the experiment were performed with identical results. (f) Interaction between Gα1^*quadD*^ and Gα1^*quadA*^ (in 77-nEYFP-N1) with full-length SymRKα or N-terminal region of SymRKα (in 78-cEYFP-N1) proteins using BiFC assay. Agrobacteria containing different combinations of Gα1 and SymRKα were infiltrated in tobacco leaves and reconstitution of YFP fluorescence due to protein-protein interaction was visualized. In all plant-based interactions (BiFC), at least five independent infiltrations were performed for each protein combination with similar results (b, d and f). Bar = 50 μm.

To confirm that the Gα1^*quadD*^ is not completely misfolded or mis-localized, or has lost its ability to interact with any protein due to multiple mutations, we tested its interaction with RGS2 NFR1α and SymRKα proteins, with which it is known to interact (Roy Choudhury & Pandey, 2015). In both yeast-based assays and *in planta* BiFC assays, the native and Gα1^*quadD*^ proteins exhibited similar interactions with RGS2, NFR1α or SymRKα proteins (Fig. 6c-f). These results confirm that the phosphorylated Gα proteins specifically fail to interact with the Gβγ dimers. Gα1, Gα1^*quadA*^ and Gα1^*quadD*^ also showed similar localization upon transfection in tobacco leaves (Fig. S8).

### Phosphorylation-based regulation of G-protein signaling during nodule formation in soybean

The data presented in this manuscript combined with our previous results led us to propose the following model for the phosphorylation-based regulation of G-protein signaling during nodule formation in soybean (Fig. 7). Gα and Gβγ proteins are negative and positive regulators, respectively, of nodule formation in soybean (Roy Choudhury & Pandey, 2013; Roy Choudhury & Pandey, 2015). During nodulation, Nod factor perception by NFR1 proteins results in receptor activation and beginning of the signal transduction (Limpens *et al.*, 2003; Madsen *et al.*, 2003; Indrasumunar *et al.*, 2011). Active NFR1 proteins phosphorylate RGS proteins, which maintain the Gα proteins in their GDP-bound, inactive form *i.e.* suppression of the negative regulators (Roy Choudhury & Pandey, 2015). Although, the GDP-bound Gα proteins are expected to remain in the trimeric form (given their high affinity for the Gβγ dimers), their phosphorylation by SymRK abolishes this binding. This would potentially allow for the constitutive signaling by Gβγ dimers. This model therefore suggests a dual control of G-protein signaling during nodulation by a pair of RLKs, which make the negative regulator (Gα) inactive and the positive regulators (Gβγ) freely available, likely resulting in a precise control of signaling during nodulation.

**Fig. 7.**
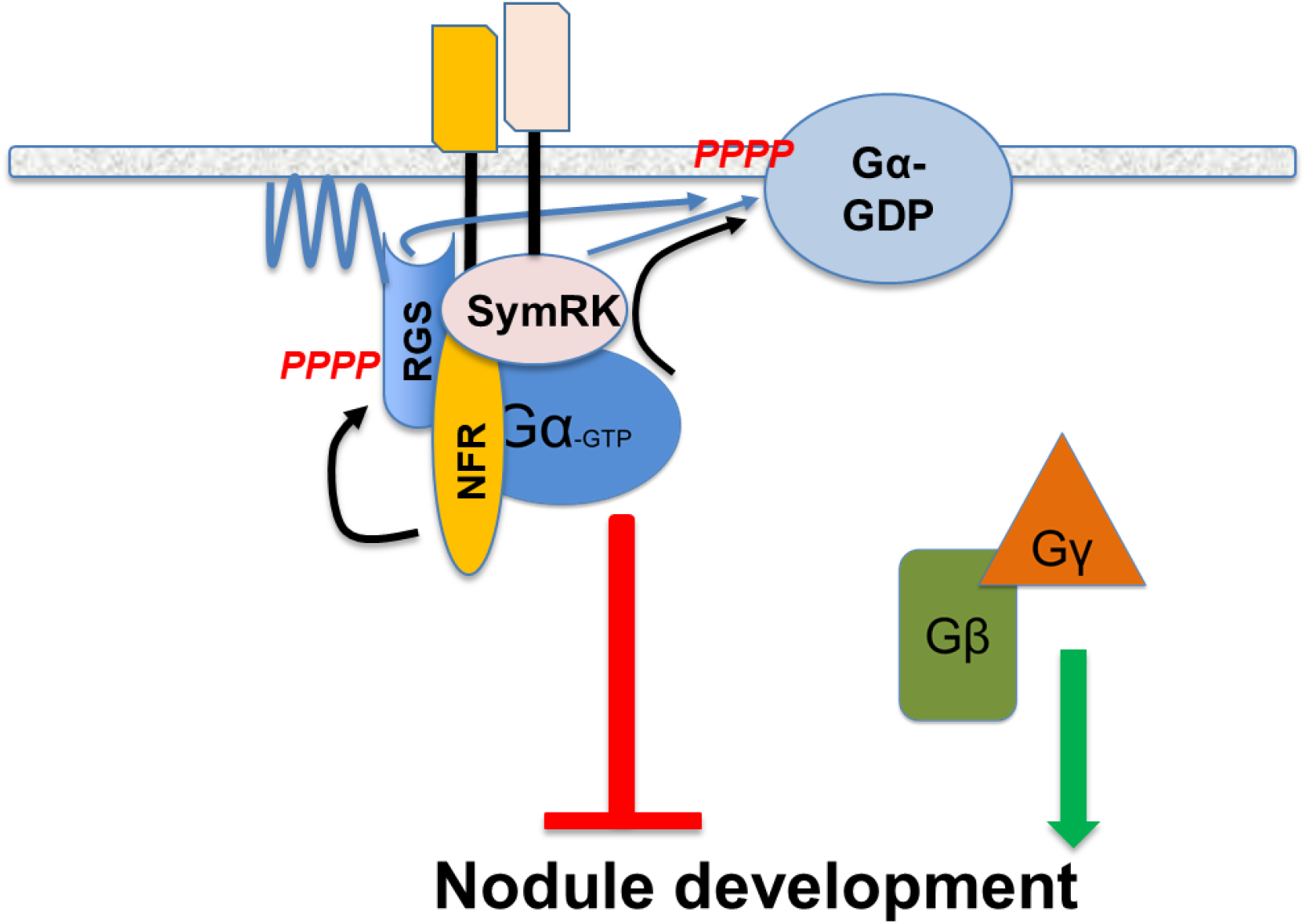
Proposed model for G-protein regulated nodule formation in soybean. A complex of RLKs regulate signaling during nodulation. NFR1 phosphorylates RGS proteins to help maintain Gα proteins in inactive conformation. SymRK phosphorylates Gα proteins to inhibit its GTP binding. Phosphorylated Gα cannot interact with Gβγ and therefore allows for constitutive signaling from free Gβγ. G-protein independent pathways, which may also exist, are not shown.

## DISCUSSION

G-proteins are key regulators of almost all aspects of growth and development in plants (Pandey & Vijayakumar, 2018; Pandey, 2019). While the details of the pathways regulated by G-proteins have been the focus of many studies in the past decades, little is known about the specific receptors, effectors, or downstream proteins, which function with G-proteins in plants. Nonetheless, it has become clear that activation/deactivation mechanisms of G-protein signaling in plants may be appreciably different from that of metazoans (Urano & Jones, 2014; Roy Choudhury & Pandey, 2016; Trusov & Botella, 2016; Tunc-Ozdemir *et al.*, 2016; Liang *et al.*, 2018; Peng *et al.*, 2018).

One critical missing piece in plant G-protein signaling is the identity of receptors that interact with G-proteins and affect their activity. Previous studies have shown that plant G-proteins interact with proteins that may have features similar to canonical GPCRs (Pandey & Assmann, 2004; Gookin *et al.*, 2008; Pandey *et al.*, 2009) however, there is no evidence yet if these GPCR like proteins have GEF activity, or if that is required for G-protein activation. Alternatively, there is growing evidence that G-proteins interact with plant specific RLKs during regulation of a variety of signaling and developmental events (Aranda-Sicilia *et al.*, 2015; Roy Choudhury & Pandey, 2015; Roy Choudhury & Pandey, 2016; Tunc-Ozdemir & Jones, 2017; Peng *et al.*, 2018). During shoot apical meristem development in maize, the Gα protein interacts with the CLV signaling pathway, although the exact mechanism of G-protein activation is not known in this regulation (Bommert *et al.*, 2013; Ishida *et al.*, 2014; Je *et al.*, 2018). On the contrary, recent evidence suggests that a phosphorylation-based mechanism operates during the FLS2 mediated defense-signaling pathway, controlling active versus inactive G-protein cycle (Liang *et al.*, 2016; *Liang et al., 2018*). In this model, the G-protein heterotrimer together with its regulatory RGS protein forms a complex with the FLS2/BAK1 receptors and a cytoplasmic kinase BIK1. Activation of FLS2 causes BIK1 activation, which phosphorylates the RGS protein, which causes release and activation of the Gα and Gβγ proteins for downstream signal transduction (Liang *et al.*, 2018). However, in this model too, the control of G-proteins by a cognate receptor is indirect, via the regulation of RGS proteins. Our previous work also demonstrated indirect regulation of G-protein cycle via NFR1-mediated RGS phosphorylation (Roy Choudhury & Pandey, 2015). Our current results corroborate this model and show that signaling during nodulation is also controlled by a direct SymRK-mediated phosphorylation of the Gα proteins.

The interaction of Gα proteins with SymRK identifies a new link between the SymRK-dependent and the G-protein-dependent signaling during nodulation (Fig. 1). The proteins also seem to affect common transcriptional targets (Fig. 5). SymRKs are known to control nodulation; however, their ligands, downstream targets or mechanistic details are not well established (Endre *et al.*, 2002; Stracke *et al.*, 2002; Capoen *et al.*, 2005; Markmann *et al.*, 2008). The SymRK interacting proteins vary from metabolic enzymes such as 3-hydroxy-3-methylglutaryl CoA reductase1 (HMGR1) to ubiquitin ligases and MAP kinases (Kevei *et al.*, 2007; Zhu *et al.*, 2008; Lefebvre *et al.*, 2010; Chen *et al.*, 2012; Den Herder *et al.*, 2012; Yuan *et al.*, 2012; Venkateshwaran *et al.*, 2015; Vernie *et al.*, 2016). Many of these especially MAP kinases and ubiquitin ligases are also well-established downstream components of G-protein signaling, and may define novel mechanistic links between these pathways. Furthermore, SymRK homologs are present in all plants, including Arabidopsis and examining their involvement in processes other than symbiosis in the context of G-protein-dependent regulation would certainly be informative.

Intriguingly, SymRKs phosphorylate Gα proteins at specific amino acids including two in their GTP-binding region (Fig. 2). Therefore, it was not surprising to find that any alteration in this region by phosphorylation (or by using a phosphomimic mutation) would abolish its GTP-binding. Assessment of the GTP-biding and GTPase activities of variant proteins that had mutations in their GTP-binding region confirmed that these are indeed unable to bind (and consequently) hydrolyze GTP (Fig. 3).

This observation is important both in the context of the regulation of signaling during nodulation proposed by our previous model (Roy Choudhury & Pandey, 2015), as well as during plant G-protein signaling in general. We had proposed that the RGS-phosphorylation by NFR1 proteins promotes GTPase activity of Gα, thereby favoring its inactive conformation (Roy Choudhury & Pandey, 2015). While this had explained the negative regulation of nodule formation by the Gα proteins, its effect on the existence of, and regulation by the Gβγ proteins remained confounding. Our data clearly showed that the Gβγ proteins are positive regulators of nodule formation (Roy Choudhury & Pandey, 2015). Because an inactive Gα would promote the formation of heterotrimer, thereby limiting the availability of free Gβγ, it wasn’t clear if the effects of Gβγ overexpression were purely due to their higher levels, and therefore relatively higher availability in free form despite high heterotrimer formation, or there were additional mechanisms involved. We had also proposed the possibility of independent regulation by Gβγ (Roy Choudhury & Pandey, 2015). The inability of the phosphorylated Gα to interact with Gβγ dimer addresses this scenario. Our current model shows that despite the presence of inactive Gα proteins, the Gβγ dimer is free for constitutive signaling and therefore can positively regulate the downstream signaling pathway (Fig. 7).

Some metazoan examples have also found Gα phosphorylation and resultant reduced affinity of phosphorylated Gα for the Gβγ dimer (Fields & Casey, 1995; Kozasa & Gilman, 1996). This suggests that not only the activation/deactivation of the G-protein cycle but also the relative availability of the subunits may be a general, but yet unexplored regulatory mechanism. One of the serine residues present in the GTP-binding region, which is a target for phosphorylation, is conserved in all Gα proteins examined. Even though we have explored the effects of this phosphorylation in the context of soybean nodulation, given the involvement of G-proteins in almost all aspects of plant growth and development, it might have additional, broadly applicable consequences.

Another intriguing aspect of this regulation is the seemingly opposite roles of Gα versus Gβγ proteins in controlling signaling. As per the metazoan paradigm, in most cases, the Gα protein is the active signaling entity and the role of Gβγ dimer is to secure it in an inactive conformation. In plants, it may be the opposite, where at least in some cases the Gβγ proteins are the active regulators and the role of the Gα proteins appears to be to keep them in inactive heterotrimeric conformation, unless activated by mechanisms such as receptor-mediated Gα phosphorylation. While we do not know the general applicability of such a mechanism yet, given the significant differences that exist in plant versus metazoan G-protein signaling regulation, it may be widespread. Such a regulation could also be important in the context of the extra-large Gα (XLG) proteins and the availability of Gβγ proteins in plants. Plant genomes encode canonical as well as XLG proteins and in species such as Arabidopsis, maize and rice, one canonical and 3-5 extra large Gα proteins share a single Gβ protein (Chakravorty *et al.*, 2015; Maruta *et al.*, 2015; Pandey & Vijayakumar, 2018; Wu *et al.*, 2018). The inability of a phosphorylated Gα to interact with Gβγ may change the stoichiometry of XLG-Gβ interactions. Interestingly the serine present in the GTP-binding pocket of Gα is also conserved in XLG proteins, predicting that these may also be phosphorylated, and potentially regulated by the activity of receptor-like (or other) kinases. The role of XLG proteins has not been explored during regulation of nodulation (there are more than 10 copies in soybean), but they are key regulators of defense signaling and function in parallel with the canonical Gα proteins in Arabidopsis. Additional research focused in simpler systems *e.g.* Arabidopsis or *Medicago* will help address some of these points.

Our model portrays an example of the complex regulatory mechanisms that exist in plant signaling and development and uncovers a novel signaling module, which links signal perception by the plasma membrane-localized receptors to their immediate downstream targets during nodulation. Nodulation is an extremely energy-demanding process and plants carefully control nodule development based on the nitrogen availability, environment conditions and overall growth (Mortier *et al.*, 2012; Downie, 2014; Nishida & Suzaki, 2018). In addition to the inbuilt mechanisms such as autoregulation of nodulation (AON) pathway, which incidentally are also controlled by RLKs (Krusell *et al.*, 2002; Nishimura *et al.*, 2002; Searle *et al.*, 2003; *Schnabel et al.*, 2005; Crook *et al.*, 2016), there may exists several parallel pathways, and G-proteins likely represent one particular regulatory module. Furthermore, given the diverse range of responses regulated by G-proteins, and the multitude of RLKs they interact with, similar mechanisms may be operative during additional growth and developmental pathways. A clearer understanding of such regulatory mechanisms will allow for their precise manipulation, resulting in crops that are more efficient, for the future needs.

## AUTHOR CONTRIBUTIONS

SP and SRC conceived the project. SRC performed all experiments with input from SP. SRC and SP wrote the manuscript.

## ACKNOWLEDGMENTS

The authors sincerely thank Dr. Lucia Strader (Washington University, St. Louis) for access to her microscope and Dr. Elizabeth Kellogg (Danforth Center) for useful comments on the manuscript. We thank two extremely patient and diligent technicians in the lab, Laryssa Hovis and Veronica Lee for counting thousands of soybean nodules. We also acknowledge the help from Danforth Center Proteomics and Mass Spectrometry Facility and Dr. Sophie Alvarez (UNL, Lincoln) for help with phospho-peptide identification. This research is supported by NIFA/AFRI (2015-67013-22964) grant to S.P.

