## Supplemental data for "A receptor-like kinase mediated phosphorylation of Gα protein affects signaling during nodulation"

**(a)**

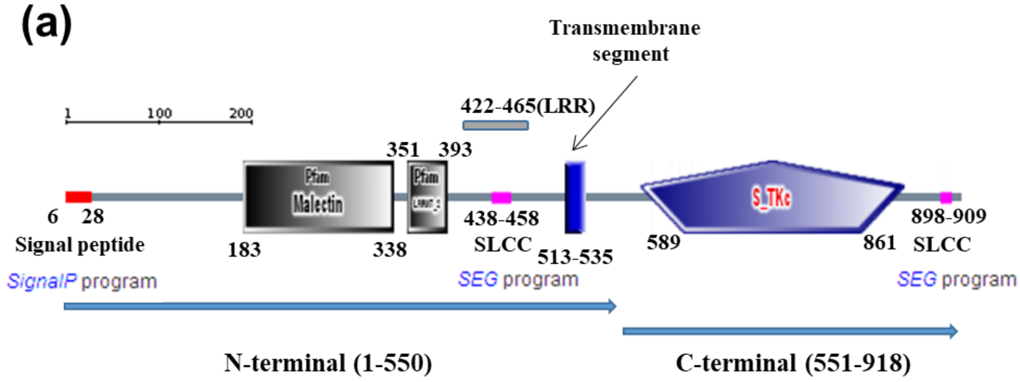


**(b)**

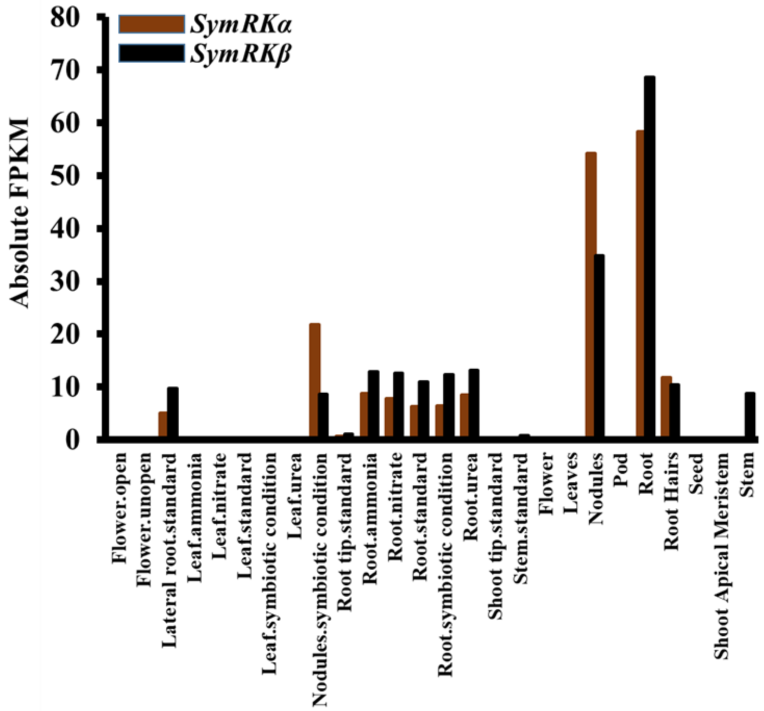


**Fig. S1. (a) Domain architecture of soybean SymRK proteins.** Soybean SymRK proteins have a signal peptide, a malectin-like domain (MLD), leucine-rich repeat (LRR) at their N-terminal region. The single transmembrane domain connects it to the C terminal region, which contains a serine-threonine kinase domain. **(b) Relative expression of *SymRKα* (**_Glyma.09G202300.1_**) and *SymRKβ* (**_Glyma.01G020100.1_**) genes in different tissues of soybean.** The absolute FPKM values of *SymRKα* and *SymRKβ* genes in different vegetative and reproductive tissues of soybean were determined from soybean RNA-seq data at Phytozome version 12.

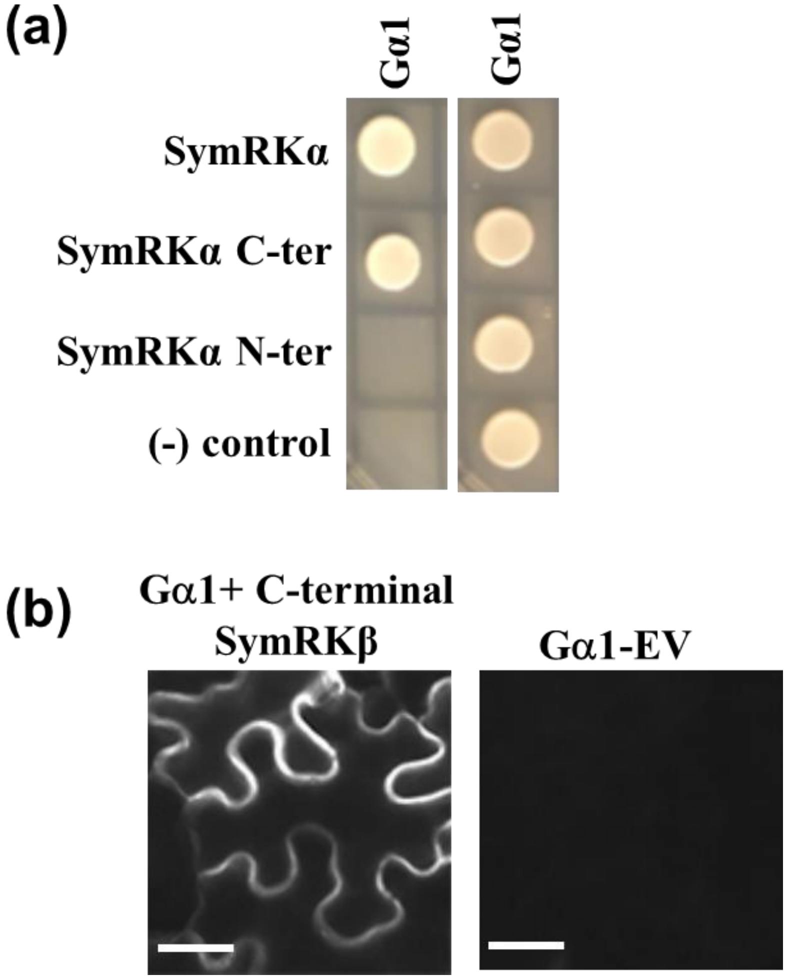


**Fig. S2. (a) Interaction between soybean Gα1 protein with full-length or N-terminal and C-terminal SymRKα proteins using split ubiquitin-based interaction assay.** In all cases, Gα1 proteins were used as NUb fusions. SymRKα proteins were used as CUb fusions. NUb-vector fusion constructs were used as negative control. Yeast were grown on selective (−Leu, −Trp) medium (*right side panel*) to verify the presence of both interacting plasmids and a more restrictive (−Leu, −Trp, −His, −Ade) medium (*left side panel*) to test for protein interactions. Two biological replicates of the experiment were performed with identical results. **(b) Interaction between Gα proteins (in 77-nEYFP-N1) with C-terminal SymRKβ or EV (in 78-cEYFP-N1) using bimolecular fluorescence complementation (BiFC) assay.** Agrobacteria containing 77-nEYFP Gα were infiltrated with 78-cEYFP SymRKβ (C-terminal) or EV in tobacco leaves. The reconstitution of YFP fluorescence due to protein-protein interaction was visualized under Nikon Eclipse E800 microscope with epi-fluorescence modules. Interaction between Gα1 and EV was used as negative control. At least five independent infiltrations were performed for each protein combination with similar results. Bar = 50 μm.

P0SGA1SWISS-PROT:P0SGA1|SGA1_XXXSGA1 (100%), 44,768.7 Da
P0SGA1SWISS-PROT:P0SGA1|SGA1_XXXSGA1
28 exclusive unique peptides, 46 exclusive unique spectra, 401 total spectra, 259/384 amino acids (67% coverage)

| | | | | | | |
|---|---|---|---|---|---|---|
| MGLVCSRSRR | FREADAEENA | QDAEIERRIE | LETKAEKHIQ | KLLLLGAGES | GKSTIFKQIK | LLFQTGFDEA |
| ELKSYIPVVH | ANVYQTIKVL | HDGSKELAQN | DFDSSKYVIS | NENQDIGEKL | SEYGGRLDYP | RLTKELAQEI |
| ETLWEDAAIQ | ETYARGNELQ | VPDCAHYFME | NLERLSDANY | VPTKEDVLYA | RVRTTGVVEI | QFSPVGENKR |
| SGEVYRLFDV | GGQRNERRKW | IHLFEGVTAV | IFCAAISEYD | QTLYEDENKN | RMMETKELFE | WVLRQPCFEK |
| TSFMLFLNKF | DIFEKKVLNV | PLNVCEWFKD | YQPVSTGKQE | IEHAYEFVKK | KFEELYFQST | APDCVDRVFK |
| IYQATALDQK | LVKKTFKLVD | ETLRRRNLFE | AGLL | | | |

| Peptide sequence | Peptide identification probability | Mascot Ion score | Modifications identified by spectrum |
|---|---|---|---|
| DYQPVSTGKQEIEHAYEFVK | 99.70% | 26.7 | Phospho (+80) |
| DYQPVSTGKQEIEHAYEFVK | 99.70% | 26.4 | Phospho (+80) |
| LLLLGAGESGKSTIFK | 99.70% | 17.5 | Phospho (+80) |
| LLLLGAGESGKSTIFK | 99.70% | 17.3 | Phospho (+80) |
| LLLLGAGESGKSTIFK | 99.70% | 17.3 | Phospho (+80) |
| LLLLGAGESGKSTIFK | 99.70% | 23.2 | Phospho (+80) |
| LLLLGAGESGKSTIFK | 99.70% | 55.8 | Phospho (+80) |
| LLLLGAGESGKSTIFK | 99.70% | 65.1 | Phospho (+80) |
| YVISNENQDIGEKLSEIGGR | 99.70% | 75 | Phospho (+80) |
| YVISNENQDIGEKLSEIGGR | 99.70% | 43 | Phospho (+80) |

**Fig. S3. Detection of phosphorylated amino acid residues in Gα1 by LC-MS/MS after *in vitro* phosphorylation assay.** The figure shows the Gα1 protein sequence identified with 67% coverage (highlighted in yellow) using a peptide probability threshold of 99% calculated according to the Peptide Prophet algorithm. Phosphorylation modifications of serine and threonine residues were identified as indicated in the figure by brown circles. Other experimental modifications (oxidation of methionine and carbamidomethylation of cysteine) are also shown in the figure although not relevant here. The localization probability of the phosphorylation sites was further tested using Scaffold PTM (v2.1.2.1 Proteome Software). The threshold was set to 99% localization probability. The lower panel shows Mascot ion score.

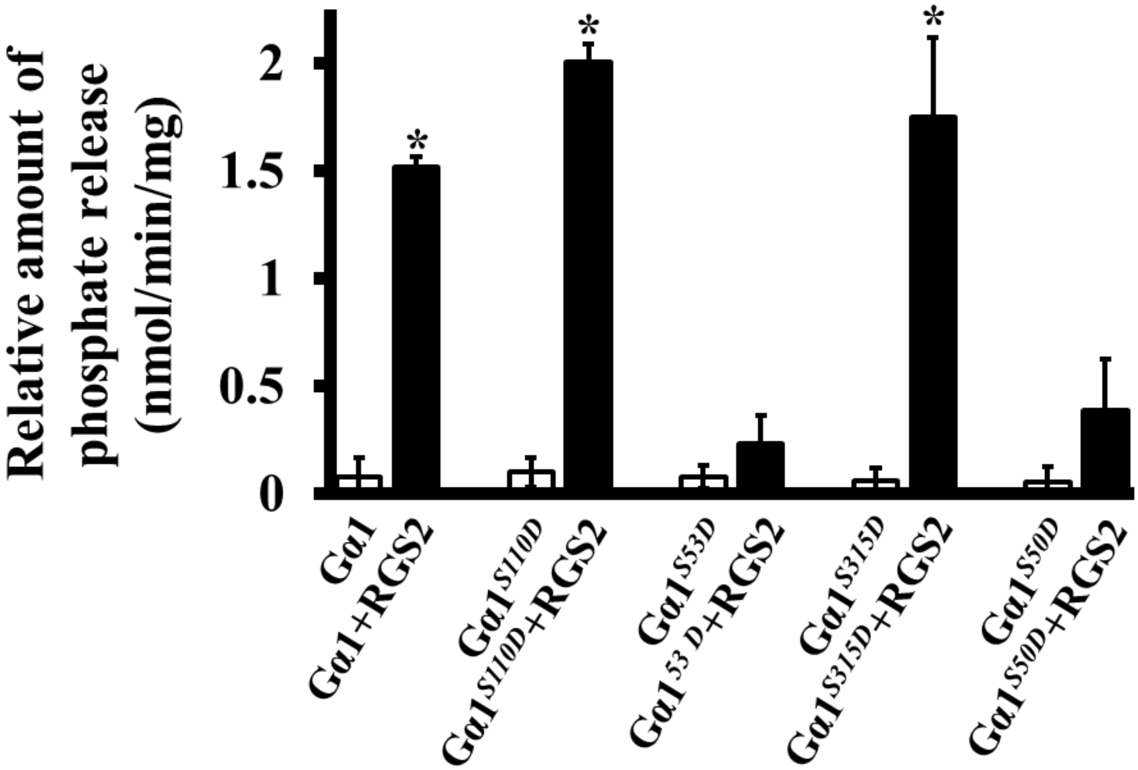


**Fig. S4. Effect of RGS2 on native and phospho-mimetic Gα1s.** Rate of Pi release due to the GTPase activity of native and phospho-mimic (Gα1$^{S110D}$, Gα1$^{S53D}$, Gα1$^{S315D}$, Gα1$^{S50D}$) versions of Gα1, inherently or in the presence of RGS2 proteins. Experiments were repeated three times, and data were averaged. Error bars represent the mean ±S.E.

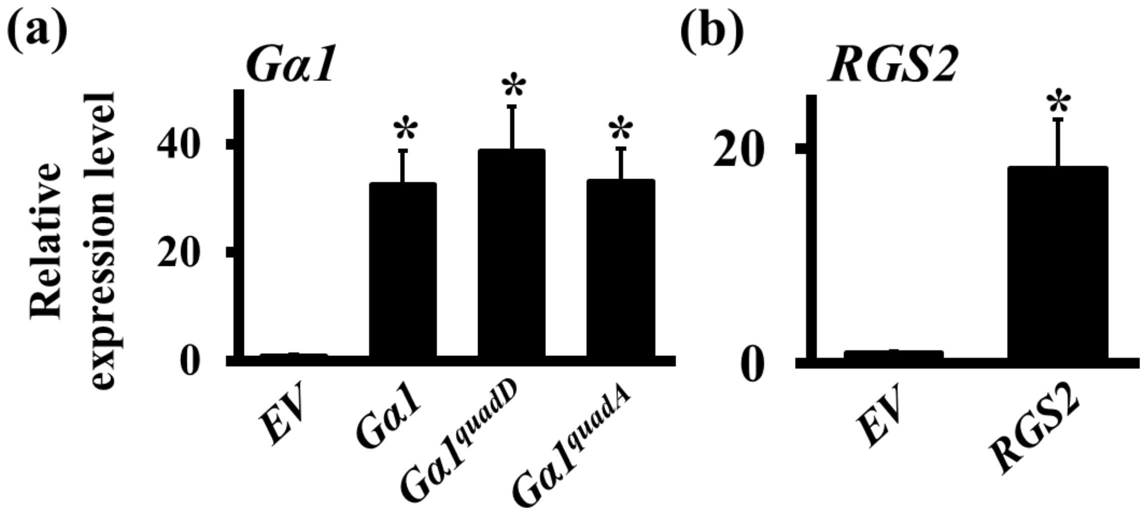


**Fig. S5. Expression levels of *Gα1* and *RGS2* genes in *Gα1* and *RGS2*-overexpressing transgenic hairy roots. (a)** Hairy roots of soybean were collected from native, phospho-dead (*Gα1*$^{quadA}$) and phospho-mimetic (*Gα1*$^{quadD}$) versions of *Gα1* over-expression lines at 4 dpi with *B. japonicum*. Fold change of gene expression was determined by comparing the transcript levels of *Gα1* gene in overexpression lines to their expression in *EV* control lines by real-time quantitative PCR. **(b)** Hairy roots of soybean were collected from *RGS2* over-expression lines at 4 dpi with *B. japonicum*. Fold change of gene expression was determined by comparing the transcript levels of *RGS2* gene in overexpression lines to their expression in *EV* control lines by real-time quantitative PCR. Two biological replicates with three technical replicates each were used for expression analysis and data were averaged. The expression values across different samples are normalized against soybean *Actin* gene expression. Expression in *EV* lines was set at 1. Error bars represent ± SE. Asterisks (*) indicate statistically significant differences compared to *EV* control (* = $P < 0.05$).

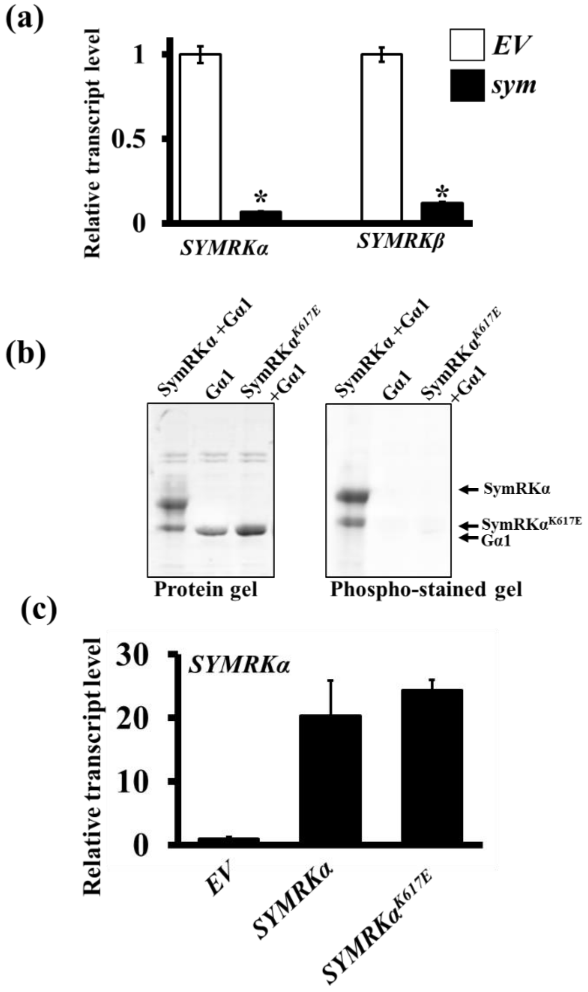


**Fig. S6. Expression levels of *SymRK* genes in *SymRK-RNAi* silenced, native and mutant *SymRKα*-overexpressing transgenic hairy roots. (a)** Hairy roots were collected from *SymRK-RNAi* lines at 4 dpi with *B. japonicum*. Expression of *SymRKα* and *SymRKβ* genes were measured by real-time quantitative PCR. **(b)** Mutant SymRKαK$^{617E}$ is inactive; it does not exhibit autophosphorylation activity and does not phosphorylate Gα protein. The mutant protein also migrates faster on gels compared to the native version. **(c)** Hairy roots were collected from native and mutant SymRK-overexpressing transgenic plants at 4 dpi with *B. japonicum*. Expression of *SymRKα* gene was measured by real-time quantitative PCR. Fold change represents expression of genes in transgenic lines in comparison to their expression in *EV* containing roots, which was set as 1. Two biological replicates with three technical replicates each were used for expression analysis and data were averaged. The expression values across different samples are normalized against soybean *Actin* gene expression. Error bars represent ± SE. Asterisks (*) indicate statistically significant differences compared to *EV* control (* = $P < 0.05$).

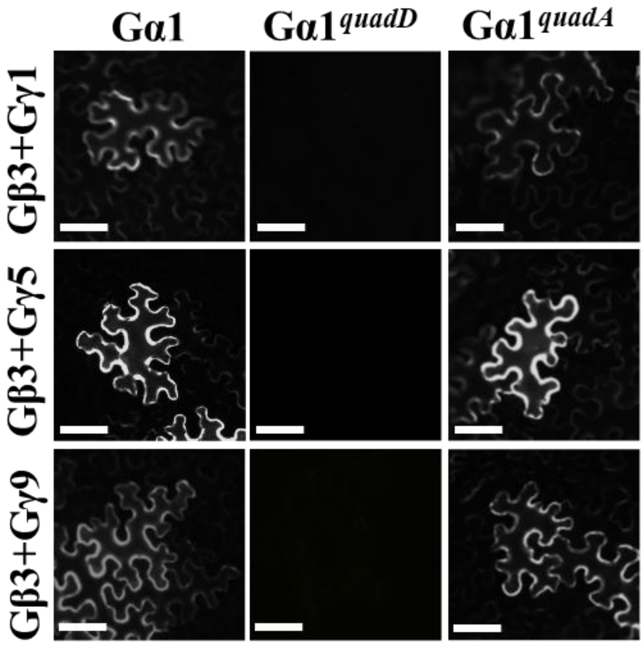


**Fig. S7. Phospho-mimic Gα cannot interact with Gβγ subunits.** Interaction between native, phospho-dead (Gα1$^{quadA}$) and phospho-mimic (Gα1$^{quadD}$) versions of Gα1 (in 77-nEYFP-N1) with Gβ3 (in 78-cEYFP-N1) and Gγ (1 or 5 or 9) proteins using BiFC assay. Agrobacteria containing different combinations of Gα1, Gβ3 and Gγ (1 or 5 or 9) subunits were infiltrated in tobacco leaves and reconstitution of YFP fluorescence due to protein-protein interaction was visualized under Nikon Eclipse E800 microscope with epi-fluorescence modules. In all plant-based interactions (BiFC), at least five independent infiltrations were performed for each protein combination with similar results. Bar = 50 µm.

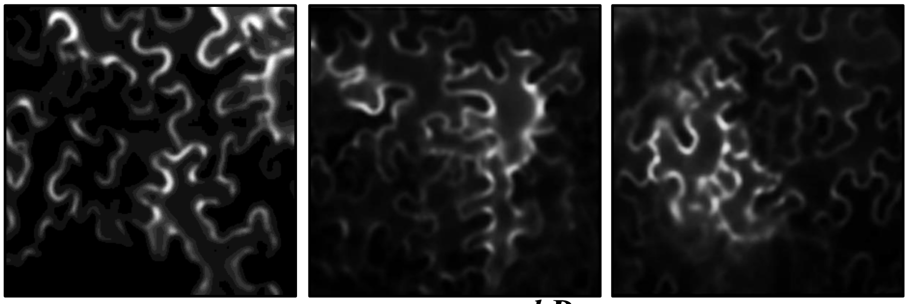


**Fig. S8. Localization of native (Gα1:YFP), phospho-mimetic (Gα1$^{quadD}$:YFP) and phospho-dead Gα1 (Gα1$^{quadA}$:YFP) in transiently transformed tobacco leaves.** At least four independent infiltrations were performed for each protein with similar results. No obvious difference in localization was observed between native and mutant protein variants.
